# The 3D Ultrastructure of C. elegans Gut Granules

**DOI:** 10.64898/2026.03.04.709038

**Authors:** Grant Archer, Sam Bartlowe, Bilegtugs Bayarbadrakh, Dawson Bok, Carolina Bushong, Anna Carroll, Megan Cole, Eric J. Dahl, Nicholas Denier, Ahmed Elewa, Hafsa El Harchi, Daniel Fisher, Hannah Grant, Rachel Grinfeld, Aliyah Grijalva, Aiden Hale, Cara Hendricks, Melek Iskandar, Sharron Kagan, Nidhi Mistry, Hope Keuneke, Lauryn Lammers, Annie Lyons, Quinn Maclin, Tatum Moore, Connor Munro, Gavin Nickerson, Brando Papalia, Kate Peacock, Lauren Ritzman, Aiden Ross, Rishi Samineni, Max Scales, Lillian Schotz, Wyatt Sikkema, Trinity Slabaugh, Autumn Sorenson, Raymond Swisher, Seyana Sutherland, Daniel Valdes, Faith Ward, Paulina Wojdyla

## Abstract

We identify an endoderm-restricted organelle in published volume electron microscopy datasets of *C. elegans* embryos. The organelle consists of a tubular ring surrounding a membrane-bound compartment harboring a prominent dense particle and exhibits a basal polarity and size concordant with canonical gut granules. This finding offers ultrastructural detail to recent evidence that gut granules are bi-lobed organelles with two distinct compartments.

## Description

The entire *C. elegans* intestine is built from endodermal cells that descend from a single blastomere [Sulston *et al*., 1983]. Restricted to intestinal cells are gut granules, lysosome-related organelles required for trace-metal storage and the synthesis of glycolipid signaling molecules known as ascarosides [Laufer *et al*., 1980, Hermann *et al*., 2005, Davis *et al*., 2009, Panda *et al*., 2017, Le *et al*., 2020]. So strict is gut granule confinement to the endoderm lineage that their ectopic appearance is evidence that a cell has aberrantly adopted an endodermal identity [Mello *et al*., 1992]. Fluorescent metabolites and birefringent crystals have endowed gut granules with striking optical properties [Cobb, 1914, Siddiqui and Babu, 1980]. The tryptophan metabolite anthranilic acid emits blue fluorescence from gut granules, and upon organismal death, floods the cytoplasm of intestinal cells in an anterior-to-posterior wave of morbid blue light [Coburn *et al*., 2013]. The source of birefringence is rhabditin, a crystallized organic substance originally identified in other nematodes and reported to fit the chemical profile of carbohydrates but apparently not metabolized upon starvation [Cobb, 1914, Laufer *et al*., 1980]. Over a century after its initial chemical characterization, the composition of rhabditin crystals remains a mystery. An intriguing property of gut granule structure emerges during zinc homeostasis. Rather than existing as a single membrane-bound compartment, gut granules appear bi-lobed, with a distinct acidified compartment and another outer compartment that expands during zinc deficiency or excess [Davis *et al*., 2009, Roh *et al*., 2012, Mendoza *et al*., 2024]. However, whether the acidified compartment is in contact with the cytosol or completely surrounded by the expansion compartment has not been resolved [Mendoza *et al*., 2024].

During an undergraduate course in cell biology, we sought the three-dimensional ultrastructure of *C. elegans* gut granules by exploring a bean-stage embryo imaged using focused ion beam scanning electron microscopy (FIB-SEM) [Santella *et al*., 2022]. We identified an organelle that is restricted to endodermal cells and has a composite structure consisting of a tubular ring encircling a membrane-bound compartment harboring a prominent, densely stained particle (**Figure 1a-c** and **Extended Data**). The identified organelle was not observed in any other cell-type in the dataset. Exclusive presence in endodermal cells was also observed in a comma-stage embryo imaged with FIB-SEM, and two later stages (1.5-fold and two-fold) imaged with array tomography [Santella *et al*., 2022] (**Figure 1d**). Additionally, the organelle in question displayed a notable asymmetric distribution consistent with the reported basal polarization of gut granules in the bean-stage [Brandt *et al*., 2022] (**Figure 1a**). We counted 314 organelles in the sixteen endodermal cells of the bean-stage embryo (median = 19, minimum = 10, maximum = 31), of which 293 included the densely stained particle (93.3%) (**Methods**). In the four aforementioned embryonic stages examined, the length of the organelle at the longest axis ranged from 0.5 to 1.4 μm with an average of 0.80 μm (s.d. = 0.15 μm, *n* = 72) (**Methods**). Earlier investigations of *C. elegans* gut granules reported a mean diameter of 0.78 μm in the embryonic pretzel-stage [Barrett and Herman, 2016]. Taken together, we have identified an organelle with a cell-type specificity, asymmetric distribution and size that all correspond to canonical gut granules. No other observed organelle shared such privilege. The tubular ring invokes the expansion compartment identified by Mendoza and colleagues, which would entail that the encircled part corresponds to the acidified compartment [Mendoza *et al*., 2024]. Finally, the densely stained particle is suggestive of rhabditin.

**Figure 1.**
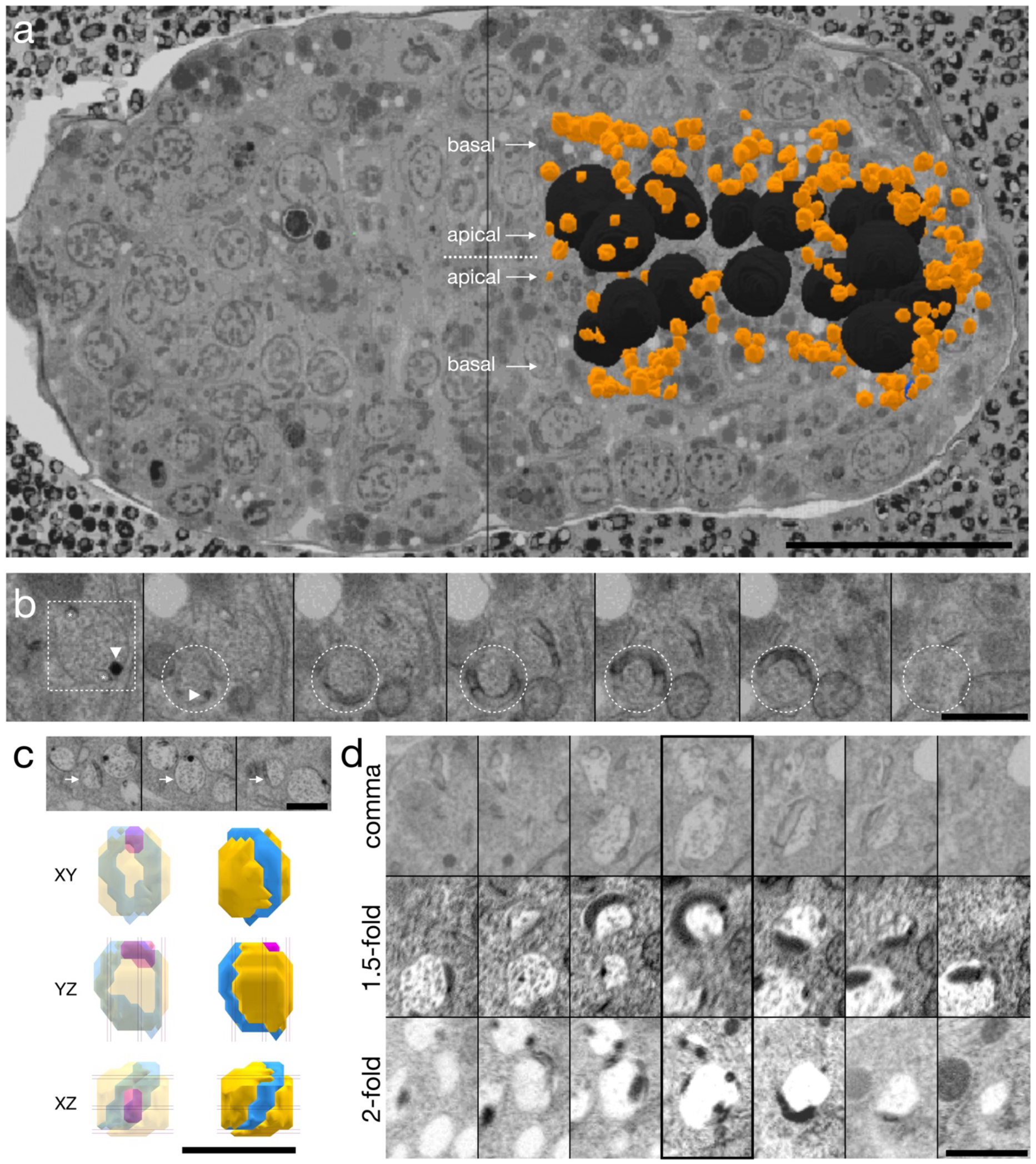
A *C. elegans* endoderm-restricted organelle. **(a)** Ventral view of annotated organelle (orange) in a bean-stage embryo against a FIB-SEM section (endodermal nuclei, black). Note the basal polarity. (b) Serial FIB-SEM sections of organelle in the bean-stage embryo. Dotted box marks an organelle; white triangle indicates a densely stained particle; asterisk denotes a tubular ring cross-section; dotted circle highlights a neighboring orthogonally oriented organelle revealing the tubular ring. (c) Four organelles in a bean-stage endodermal cell shown across three sections. Arrow indicates a reconstructed organelle below displayed in three axes: tubular ring (blue), encircled compartment (yellow), densely stained particle (magenta). Vertical and horizontal lines correspond to the EM sections above. (d) Serial sections of representative organelles in three additional embryonic stages. Each middle section (thick border) consists of two organelles. Scale bars: 10 μm (a); 1 μm (b-d).

## Methods

### Annotation, counting and measurement

Datasets were accessed through their original publication and explored using WEBKNOSSOS [Boergens *et al*., 2017]. The organelle in question was counted after annotation. Note that in the bean-stage, the cell Ealaav/d has undergone karyokinesis but not cytokinesis and had the maximum number of organelles (31 organelles). The length of the organelle along its longest axis was measured in WEBKNOSSOS for three different organelles in six different cells. This was repeated in each of the four developmental stages each in its original axis of acquisition (i.e., the XY axis for bean, 1.5-fold and two-fold stages and the YZ axis for the comma stage). Calculations were performed on all 72 measurements.

### Statistical analysis

Averages and standard deviations were calculated in Microsoft Excel.

## Extended data

https://webknossos.org/links/BLbIDFurBwDwZYqN A public link to a forked annotation of the Santella *et al*. 2022 bean-stage FIB-SEM dataset with the annotations used for **Figure 1**. Note that the names used for each cell are based on the identification retrieved in the original annotation.

## Acknowledgements

We thank Irina Kolotueva, Anthony Santella, Zhriong Bao and Norman Rzepka for early discussion of our work. Greg Hermann offered generous feedback on the project and the manuscript. This work was supported by Miami University startup grant MS43099.

## Author Contributions

ED, HEH, QM, CM, BP focused on gut cells and confirmed the presence of the organelle. SB, BB, ND, LR, RS, MS, RS focused on hypodermal cells and confirmed absence of the organelle. MC, HK, TS, DV focused on seam cells and confirmed absence of the organelle. CB, SK, LL, AL, LS, WS focused on pharyngeal cells and confirmed absence of the organelle. DF, AG, AH, CH, NM, AR, AS focused on neuronal cells and confirmed absence of the organelle. DB, HG, GN, KP, SS, FW, PW focused on muscle cells and confirmed absence of the organelle. GA, AC, RG focused on excretory cells and confirmed absence of the organelle. MI and TM focused on gonad cells and confirmed absence of the organelle. AE designed and supervised the project, performed the annotations and quantifications herein, confirmed restriction of the identified organelle to endodermal cells in all datasets examined, and wrote the manuscript.

